# Protein-Protein Interactions Shape Genomic Autoimmunity in the Adaptively-Evolving Rhino-Deadlock-Cutoff (RDC) Complex

**DOI:** 10.1101/2020.11.12.380154

**Authors:** Erin S. Kelleher

## Abstract

The piRNA pathway is a genomic defense system that controls the movement of transposable elements (TEs) through transcriptional and post-transcriptional silencing. While TE defense is critical to ensuring germline genome integrity, it is equally critical that the piRNA pathway avoids autoimmunity in the form of silencing host genes. Ongoing cycles of selection for expanded control of invading TEs, followed by selection for increased specificity to reduce impacts on host genes, are proposed to explain the frequent signatures of adaptive evolution among piRNA pathway proteins. However, empirical tests of this model remain limited, particularly with regards to selection against genomic autoimmunity.

I examined three adaptively evolving piRNA proteins, Rhino, Deadlock and Cutoff, for evidence of interspecific divergence in autoimmunity between *Drosophila melanogaster* and *D. simulans*. I tested a key prediction of the autoimmunity hypothesis: that foreign heterospecific piRNA proteins will exhibit enhanced autoimmunity, due to the absence of historical selection against off-target effects. Consistent with this prediction, full-length *D. simulans* Cutoff, as well as the *D. simulans* hinge domain of Rhino, exhibit expanded regulation of *D. melanogaster* genes. I further demonstrate that this autoimmunity is dependent on known incompatibilities between *D. simulans* proteins or domains and their interacting partners in *D. melanogaster*. My observations reveal that the same protein-protein interaction domains that are interfaces of adaptive evolution in Rhino and Cutoff also determine their potential for autoimmunity.

## INTRODUCTION

The Piwi-interacting RNA (piRNA) pathway is an RNA-mediated silencing pathway that controls the mobilization of transposable elements (TEs) in metazoan germlines (reviewed in Czech et al. 2018; Ozata et al. 2019). piRNA pathway evolution is exceptionally dynamic, including both gene duplication and rapid adaptive protein evolution (Obbard et al. 2009; Kolaczkowski et al. 2011; Simkin et al. 2013; Yi et al. 2014; Lewis et al. 2016; Palmer et al. 2018; Crysnanto & Obbard 2019). This adaptive evolution is likely required to maintain control of genomic TEs, which change rapidly in presence and abundance over short evolutionary time periods (Kidwell 1983; Naito et al. 2006; Yang & Barbash 2008; El Baidouri & Panaud 2013; Gilbert et al. 2010; Reiss et al. 2019). However, it is equally crucial that the piRNA pathway avoids collateral damage in the form of off-target silencing of host genes. piRNA pathway evolution, therefore, is proposed to reflect a trade-off between maximizing TE regulation while minimizing genomic autoimmunity (Blumenstiel et al. 2016).

The piRNA pathway has been most extensively characterized in *Drosophila melanogaster* and consists of >30 proteins with diverse functional roles in piRNA transcription, maturation and enforcement of transcriptional and post-transcriptional silencing (reviewed in Senti & Brennecke 2010; Ozata et al. 2019). Proteins that establish piRNA transcription play potentially critical roles in both adaptation to genomic TEs and avoidance of autoimmunity by determining the repertoire of cellular piRNAs (Blumenstiel et al. 2016; Palmer et al. 2018). Indeed, three key regulators of piRNA precursor transcription: Rhino (Rhi), Deadlock (Del) and Cutoff (Cuff) are among the most adaptively evolving piRNA proteins in the genus *Drosophila* (Figure 1A; Vermaak et al. 2005; Simkin et al. 2013; Blumenstiel et al. 2016; Palmer et al. 2018). Rhi recognizes piRNA-producing loci known as piRNA clusters, and together with Del and other co-factors, recruits RNA-polymerase II to initiate precursor transcription (Andersen et al. 2017). The Rhino-Deadlock-Cutoff (RDC) complex further suppresses mRNA transcription at piRNA clusters, and ensures the transport of precursor transcripts to cytoplasmic sites of piRNA maturation (Mohn et al. 2014; Zhang et al. 2014; Chen et al. 2016). The RDC complex could therefore exhibit autoimmunity through suppressing mRNA transcription or misappropriating genic mRNAs to piRNA processing bodies.

**Figure 1.**
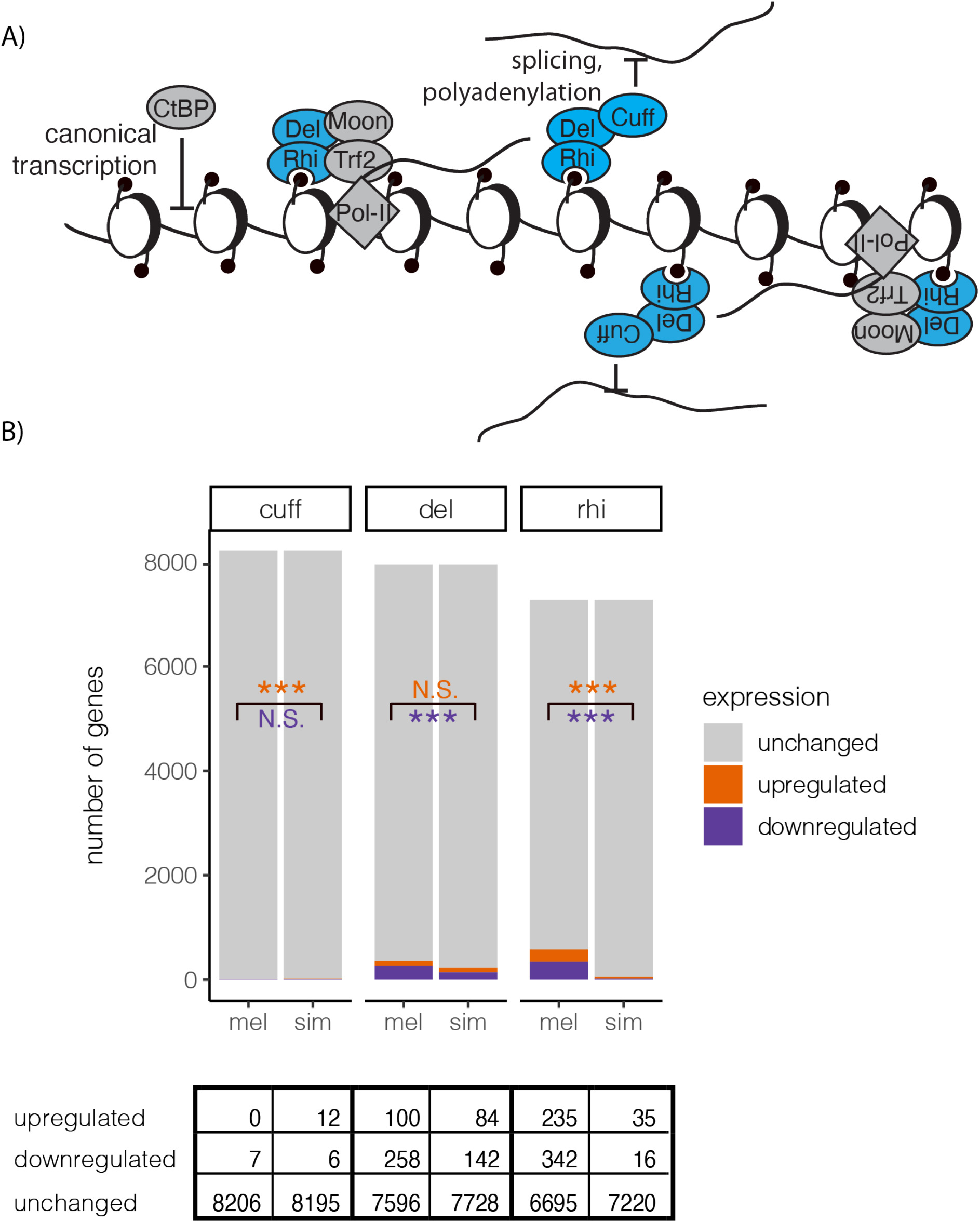
RDC regulation of piRNA precursors and host genes. A) Schematic of known Cuff, Rhi and Del functions in piRNA precursor transcription. Rhi and Del, together with Moon and Trf2 act to recruit bidirectional transcription of piRNA clusters (Andersen et al. 2017). Rhino, Del and Cuff further specify piRNA precursors through suppressing splicing, polyadenylation and termination (Mohn et al. 2014; Zhang et al. 2014; Chen et al. 2016). CtBP represses canonical, promoter-dependent transcription at piRNA clusters (Parhad et al. 2020). B) Genes that are upregulated, downregulated, and unchanged by *cuff, del* and *rhi* transgenic rescues as compared to the corresponding mutant background are indicated. Results of Fisher’s exact test (*cuff*, upregulated) or ***X***^2^ test-of-independence (all others, *df*=1) indicating differences in the proportion of genes positively (orange, top) or negatively (purple, bottom) regulated between transgenic rescues are also indicated. N.S. denotes P-value > 0.05, * denotes P-value < 0.05, *** denotes P-value < 0.001.

Interspecific complementation, in which a loss-of-function mutant is rescued by a wildtype allele from another species, provides a powerful approach for uncovering functional differences in proteins resulting from adaptive evolution (Aruna et al. 2009; Flores et al. 2015; Parhad et al. 2017; Brand et al. 2018). Using this approach, it was revealed that *D. simulans* alleles of Rhi, Del and Cuff are unable to complement *D. melanogaster* mutant backgrounds, and exhibit drastic defects in piRNA biogenesis and TE regulation (Parhad et al. 2017; Yu et al. 2018; Parhad et al. 2020). Furthermore, these functional deficits of *D. simulans* alleles result from incompatibilities with interacting cofactors in *D. melanogaster* (Parhad et al. 2017; Yu et al. 2018; Parhad et al. 2020). In the context of genomic autoimmunity, it is predicted that *D. simulans* alleles will also exhibit enhanced off-target effects in *D. melanogaster*, because there is no evolutionary history of purifying selection against targeting host mRNAs in a foreign genome. We recently uncovered this signature of expanded autoimmunity in *D. simulans* alleles of the cytoplasmic piRNA proteins Aubergine and Armitage (Wang et al. 2020). However, despite their stronger signatures of adaptive evolution and greater potential to initiate off-target effects, the autoimmunity of *D. simulans* alleles of Rhi, Del and Cuff have never been investigated.

Here, I examined the autoimmunity of *D. simulans* Rhi, Del and Cuff in a *D. melanogaster* background using published RNA-seq, small RNA-seq and ChIP seq data from interspecific complementation experiments (Parhad et al. 2017, 2020). For Rhi and Cutoff, I discovered disparate patterns of increased autoimmunity, which are determined by their incompatibilities with *D. melanogaster* cofactors. In the case of Rhi, increased autoimmunity is exhibited by the *D. simulans* hinge domain, but is masked by an incompatibility between *D. simulans* chromo shadow domain and *D. melanogaster* Del. In contrast, *D. simulans* Cuff increases the expression of hundreds of genes, potentially through the non-functional sequestration of *D. melanogaster* transcriptional regulators.

## RESULTS

### *Drosophila simulans* alleles of the RDC complex do not exhibit expanded silencing of *D. melanogaster* host genes

The genomic autoimmunity model predicts that foreign *D. simulans* alleles will exhibit expanded negative-regulation of host genes when compared to their native *D. melanogaster* counterparts (Blumenstiel et al. 2016). To test this prediction, I identified genes that were upregulated and down-regulated in ovaries by *D. melanogaster* and *D. simulans* transgenic rescues of *rhi, del* and *cuff*, when compared to an unrescued mutant background (Figure 1B, Table S1, S2). For both *del* and *rhi*, significantly fewer genes were negatively-regulated by *D. simulans* transgenes than by *D. melanogaster* transgenes (Del: ***X***^2^ = 27.27, *df* = 1, P-value = 1.77 x 10^-7^, Rhi: ***X***^2^ = 304.31, *df* = 1, P-value < 10^-15^). For *cuff*, the *D. melanogaster* and *D. simulans* transgenes negatively-regulate a similarly small number of genes; however, *D. simulans cuff* upregulates significantly more genes than *D. melanogaster cuff* (Fisher’s exact test *p* =0.0005). Collectively, these results provide no obvious support for the autoimmunity model.

In the case of *D. simulans del* and *rhi*, reduced impacts on host gene expression are potentially explained by incompatibilities with *D. melanogaster* interactors, which prevent the production of piRNAs and therefore the manifestation of autoimmunity. *Drosophila simulans* Rhi is unable to interact with *D. melanogaster* Del, which abrogates piRNA transcription (Parhad et al. 2017; Yu et al. 2018). *Drosophila simulans* Del is similarly unable to promote piRNA biogenesis in *D. melanogaster*, most likely due to incompatibilities with other cofactors (Parhad et al. 2017). Robust examination of autoimmunity of *D. simulans* proteins may therefore require restoring their capacity to interact with cofactors, so that resulting differences in gene regulation are revealed.

### Divergence in gene regulation by *D. simulans* Rhi is masked by its incompatibility with *D. melanogaster* Del

For *D. simulans* Rhi, the incompatibility with Del is caused by the chromo shadow domain (Figure 2A; Parhad et al. 2017). Chimeric transgenes combining the *D. melanogaster* chromo shadow domain with *D. simulans* domains elsewhere in the protein are functional for piRNA biogenesis and TE regulation (Figure 2B; Parhad et al. 2017). Rhi is an HP1 homolog that, in addition to the chromo shadow domain, contains a chromo and a hinge domain (Vermaak et al. 2005; reviewed in Vermaak & Malik 2009, Figure 2A). The chromo domain is responsible for binding to the histone modification H3K9me3 (Le Thomas et al. 2014; Mohn et al. 2014; Yu et al. 2015), and the hinge domain of another *D. melanogaster* HP1 homolog determines targeting to heterochromatin (Smothers & Henikoff 2001). Both the chromo and hinge domains therefore have the potential to establish off-target effects by localizing Rhino to genic regions.

**Figure 2.**
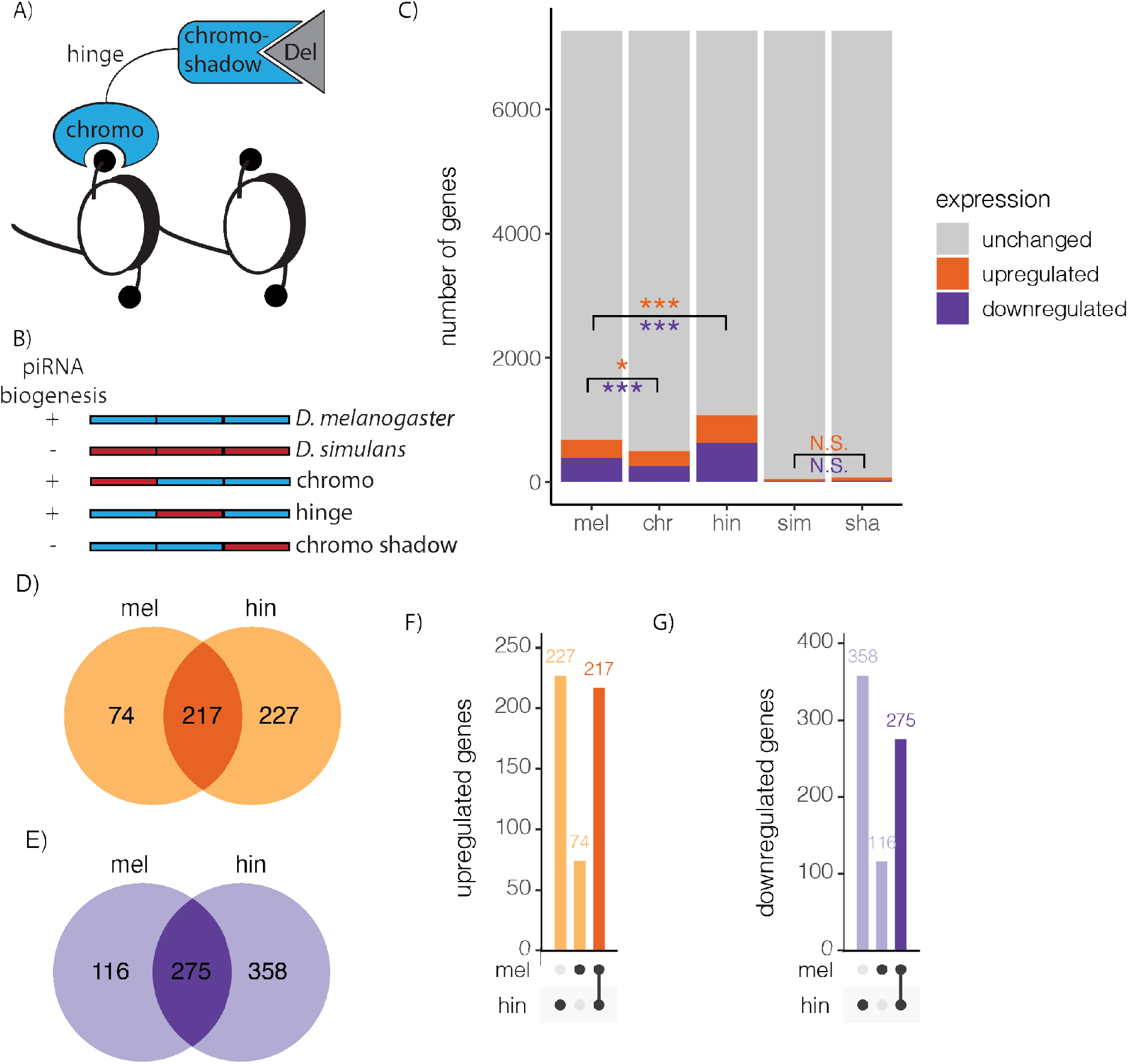
Divergence in host gene regulation individual Rhi domains. A) Cartoon of 3 Rhino domains. B) Schematic of *D. melanogaster, D. simulans* and three fusion rescue proteins from generated by Parhad *et al.* (2017). Rescue +/- indicates whether the fusion construct was previously reported to rescue piRNA biogenesis and TE regulation (Parhad et al. 2017). C) Genes that are upregulated, downregulated, and unchanged as compared to the corresponding mutant background are compared between *D. melanogaster* (mel) and *D. simulans* (sim) transgenic rescues as well as the chromo (chr), hinge (hin) and chromo shadow (sha) fusion transgenes. Results of ***X***^2^ tests-of-independence indicating differences in the proportion of genes positively (orange, top) or negatively (purple, bottom) regulated between a fusion transgene and the *D. melanogaster* or *D. simulans* transgenes are indicated. D-E) Venn diagrams and F-G) upset plots comparing genes (B and D) positively and (C and E) negatively regulated by the *D. melanogaster* (mel) and hinge (hin) transgenic rescues of *rhi* as compared to unrescued mutants. Genes that are uniquely positively regulated by one of the two transgenic constructs are shaded lighter, while those regulated by both transgenic constructs in the same direction are shaded darker. N.S. denotes P-value > 0.05, * denotes P-value < 0.05, *** denotes P-value < 0.001

By comparing the gene regulatory effects of *D. simulans, D. melanogaster* and their chimeric *rhi* transgenes to unrescued mutant backgrounds, I revealed interspecific divergence in the hinge and chromo domains (Figure 2B, 2C, Table S2). Consistent with abrogated function resulting from the Del incompatibility, the fusion transgene containing the *D. simulans* chromo shadow regulates only a small handfull of host genes, similar to the pure *D*. simulans transgene (Figure 2C). In contrast, the hinge and chromo fusion transgenes, which are compatible with *D. melanogaster* Del, exhibit increased regulation of host genes when compared to the *D. simulans* transgene (Figure 2C). In particular, the hinge fusion transgene also positively and negatively regulates more host genes than the *D. melanogaster* transgene (positive-regulation: ***X***^2^ = 34.47, *df* = 1, P-value = 4.32 x 10^-9^; negative regulation: ***X***^2^ = 61.46, *df* = 1, P-value = 4.53 x 10^-15^; Figure 2C-G). The pattern is stronger for negatively-regulated genes, with the hinge fusion transgene reducing the expression of 633 genes, 358 of which are not negatively-regulated by the *D. melanogaster* transgene (Figure 2E,G). Overall this pattern is consistent with increased autoimmunity of the *D. simulans* hinge domain in a *D. melanogaster* background.

While the expression of many genes could be indirectly affected by Rhi function (for example, through reduced DNA damage resulting from TE activity), genic sites of Rhi occupancy are more likely to represent true examples of genomic autoimmunity. I therefore used ChIP seq data from GFP-tagged Rhi proteins to identify the fraction of genes regulated by the *D. melanogaster* and hinge fusion constructs that exhibited a corresponding peak of Rhi occupancy within the gene body or up to 1 Kb upstream (Parhad et al. 2017, TableS3). Genes negatively-regulated by both transgenes were enriched for Rhi occupancy, consistent with a model in which Rhi reduces mRNA transcription (hinge: ***X***^2^ = 130.94, *df* = 1, P-value < 10^-15^, mel: ***x***^2^ = 12.579, *df* = 1, P-value = 0.00039). However, genes negatively regulated by the hinge fusion constructed are significantly more enriched for Rhi occupancy, with >18% (118/633) of negatively-regulated genes also being occupied by the hinge fusion Rhi protein, while only ~13% (51/391) of genes negatively-regulated by *D. melanogaster* Rhi are also occupied by the *D. melanogaster* Rhi protein (***X***^2^ = 5.5, *df* = 1, P-value = 0.019, Figure 3A vs 3B). In contrast, upregulated genes are not enriched for Rhi occupancy for either transgenic rescue (Figure 3C, 3D). Taken together these observations reveal that Rhi occupancy leads exclusively to negative regulation of adjacent genes, and that this activity is more pronounced with the *D. simulans* hinge domain.

**Figure 3.**
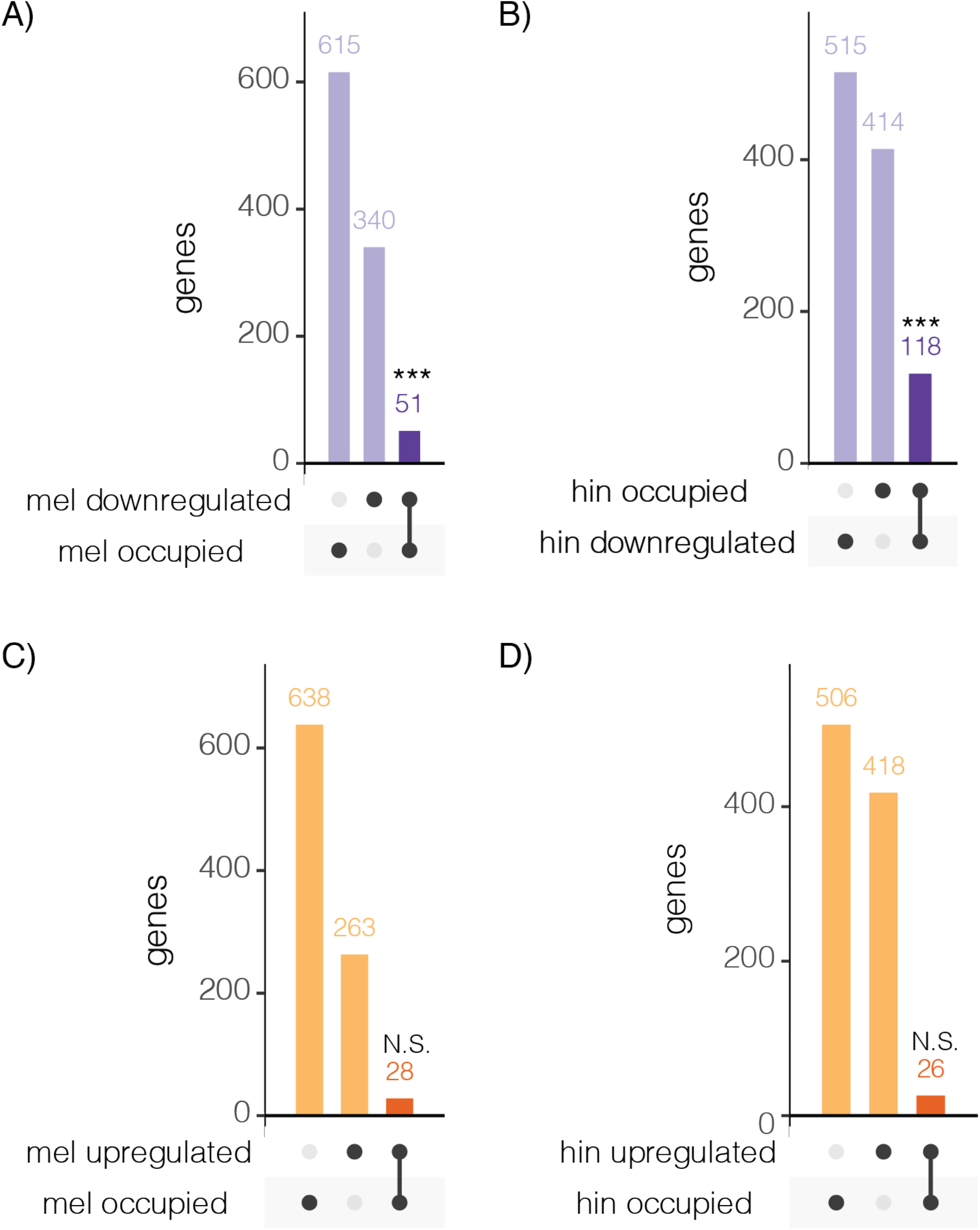
The hinge domain of *D. simulans* Rhi exhibits expanded autoimmunity of *D. melanogaster* genes. A-B) Upset plots comparing genes occupied and downregulated by *D. melanogaster* (mel, A) and hinge Rhi (hin, B). C-D) genes occupied and upregulated by *D. melanogaster* (mel, C) and hinge Rhi (hin, D). Darker colors denote an overlapping set of genes in both groups: regulated and occupied, which are the strongest candidates for autoimmunity. *** denotes P-value < 0.001 for X^2^ test-of-independence between regulation and occupancy whereas N.S. denotes P-value > 0.05.

### Expanded autoimmunity at the histone gene cluster

To better understand the expanded autoimmunity conferred by the *D. simulans* hinge domain, I examined the genes that are unique autoimmunity targets (occupied and negatively-regulated) of the hinge fusion protein when compared to the *D. melanogaster* protein (Figure 4A). The majority of novel autoimmunity targets (68 of 85) of the *D. simulans* hinge domain correspond to copies of replication-dependent histones (Figure 4A, 4B). Excluding histones, the *D. melanogaster* and hinge fusion proteins exhibit a similar number of unique autoimmunity targets (17 and 18, Figure 4A). Expanded autoimmunity of the *D. simulans* hinge domain is therefore fully-explained by its expanded regulation of histone gene copies. Interestingly, the chromo fusion construct also exhibits expanded histone autoimmunity, however, this is counterbalanced by a reduction in non-histone targets (Figure 4A, 4B). Expanded regulation of histones in a *D. melanogaster* background is therefore a shared property of both the hinge and chromo domains of *D. simulans* Rhi.

**Figure 4.**
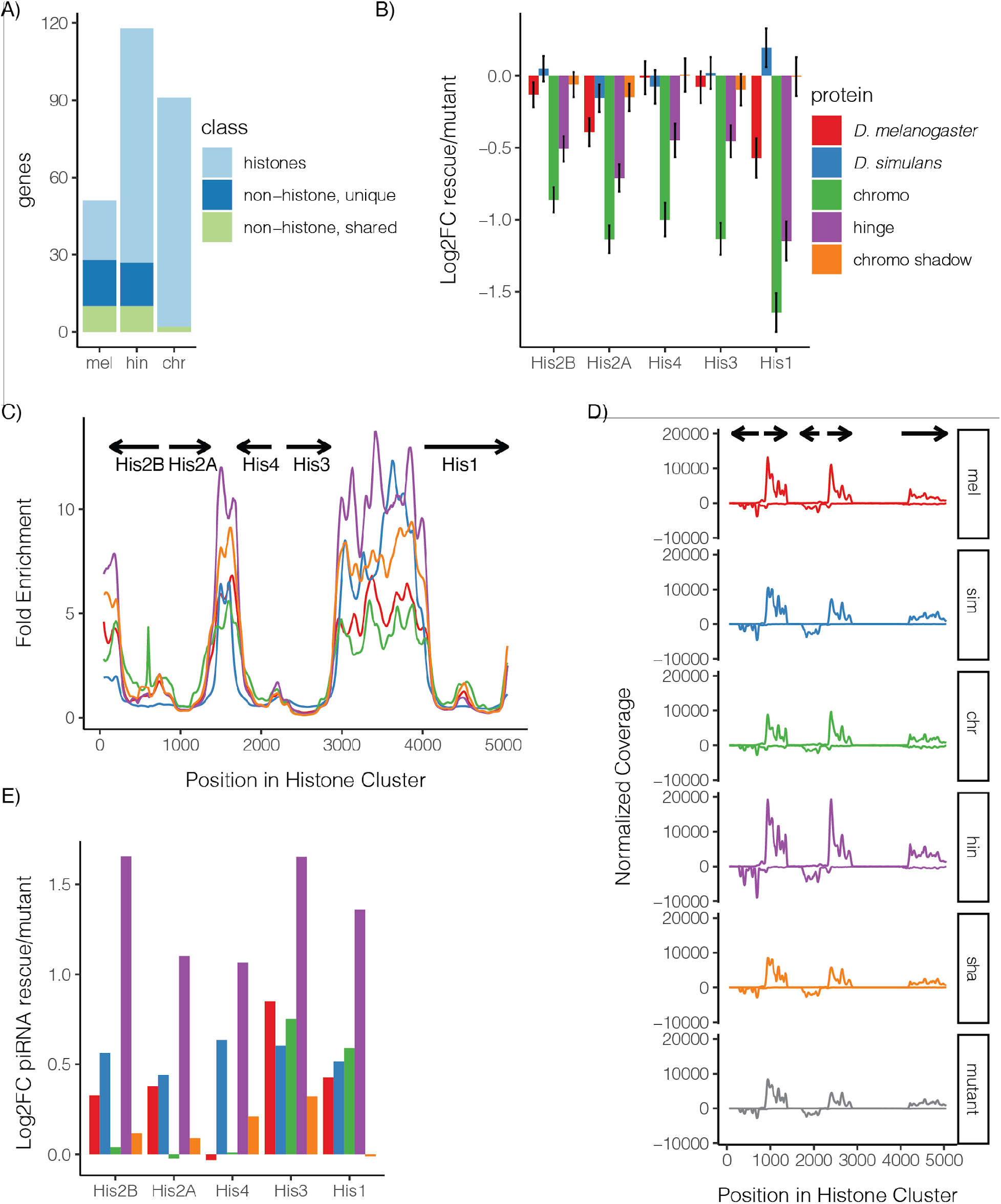
Autoimmunity at the histone gene cluster. A) Autoimmunity targets, which are both occupied and negatively regulated, are compared between the *D. melanogaster* (mel), hinge (hin) and chromo (chr) fusion proteins. B) Log2FC expression of histone copies in transgenic rescues as compared to unrescued *rhi* mutants. Due to sequence similarity between histone gene copies as well as their coordinated regulation, estimated read counts were summed across all copies to calculate differential expression. C) Sliding window of Rhi protein enrichment, as compared to input, across a single representative copy of the histone array (dm6, 2L:21,482,367-21,487,518). D) Sliding window analysis of small RNA coverage across a single representative copy of the histone array for *D. melanogaster* (mel), *D. simulans* (sim), chromo (chr), hinge (hin) and chromoshadow (sha) fusion recues, as well as the unrescued mutant. E) Log2FC piRNA abundance for each of the histone genes in transgenic rescues as compared to unrescued mutants.

Replication dependent histone genes reside in a coregulated tandem array, which includes 20-23 copies of each of the 5 histone genes. All five histones are negatively regulated (1.4-2.2 fold) by the hinge transgene, while the *D. melanogaster* transgene exhibits only modest negative regulation of *his1* and *his2A* copies (1.5 and 1.3 fold reduction, respectively; Figure 4B). Rhi-dependent differences in histone regulation could reflect differential occupancy of the histone gene cluster, or differential downstream effects of Rhi occupancy. To discriminate between these alternatives, I examined both Rhi occupancy of the histone gene cluster, and its downstream effects on histone expression and piRNA production (Figure 4B-E). To avoid complications of sequence homology among histone gene copies, these analyses were performed using a genome containing a single representative histone gene cluster as in McKay *et al.* (2015). While two occupancy peaks within the histone cluster are observable for all 5 Rhi proteins, the hinge fusion protein is the most enriched (Figure 4C), and similarly shows the greatest abundance of histone genic piRNAs (Figure 4D,E). By contrast, the *D. melanogaster* protein is significantly enriched only upstream of *his1* (although a non-significant peak 3’ to *his2A* and *his4* is observable), and exhibits only modest impacts on histone genic piRNAs. Expanded autoimmunity against the *D. melanogaster* histone gene cluster established by the *D. simulans* hinge domain is therefore associated with enhanced Rhi occupancy and downstream piRNA biogenesis.

Differences in histone gene regulation among Rhi proteins are not universally explained by differential occupancy, however. The *D. simulans* and chromo shadow fusion proteins are comparably enriched to the *D. melanogaster* protein but do not negatively regulate histone transcripts, presumably due to their inability to bind Del (Figure 4B,C). Furthermore, the chromo fusion construct exhibits negative regulation of all 5 histone genes that exceeds that of the hinge fusion construct (Figure 4A), yet the protein itself is less enriched at the histone cluster (Figure 4C). Therefore, increased negative regulation of histones established by the *D. simulans* chromo domain must occur downstream of occupancy, potentially by altering interactions between Rhi and other proteins.

### *D. simulans* Cuff regulates host genes through sequestration of CtBP

Lastly, I considered the unusual regulatory effects of *D. simulans cuff*, which exhibits expanded upregulation of host genes when compared to the *D. melanogaster* allele (Figure 1B). This expanded upregulation appears modest when compared the transgenic rescue is compared to the unrescued mutant, impacting the expression of only 12 genes (Figure 1B, 5A, 5B). However, it is revealed as quite dramatic when the gene expression profiles of the *D. simulans* and *D. melanogaster* transgenic rescues are compared to each other (Figure 5C). 159 genes are differentially expressed between the transgenic rescues, 141 of which exhibit higher expression in the presence of *D. simulans cuff*. The two transgenes exhibit opposing effects on the expression of these 141 genes, with expression values being higher in *D. simulans* rescues than mutants, but lower in *D. melanogaster* rescues (Figure 5D).

**Figure 5.**
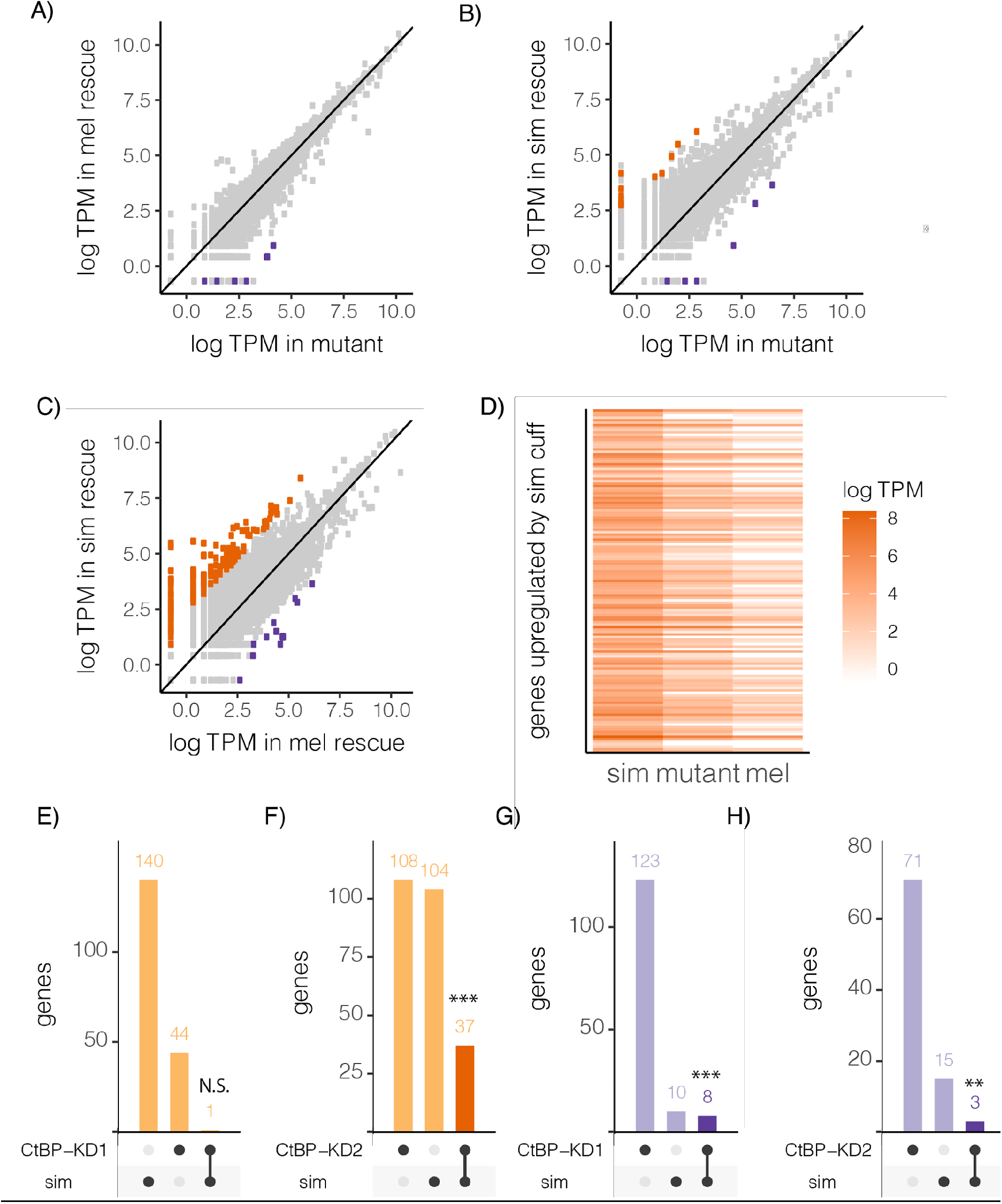
*D. simulans cuff* upregulates numerous host genes by sequestering CtBP. A-C) Correlation plots of the log-scale gene expression levels in transcripts per million (TPM) between the three *cuff* genotypes: unrescued mutant (mut), *D. melanogaster* (mel) and *D. simulans* (sim) transgenic rescues. D) Log expression levels (TPM) are compared between all three genotypes for 141 genes upregulated by the *D. simulans* (sim) as compared to the *D. melanogaster* (mel) *cuff* transgenic rescue. E-H) Upset plots comparing genes upregulated (E-F) and downregulated (G-H) by *D. simulans* as compared to *D. melanogaster* transgenic rescues and in CtBP knock-down as compared to control flies. CtBP-KD1: *P{KK108401}VIE-260B, CtBP-KD2:P{GD4268}v37609*. Results of Fisher’s exact test (CtBP-KD2, downregulated) or ***X***^2^ test-of-independence (all others, *df*=1) indicating significance of overlap in upregulated or downregulated genes between *D. simulans cuff* transgenic rescues CtBP-KD are also indicated. N.S. denotes P-value > 0.05, ** denotes P-value < 0.01, *** denotes P-value < 0.001.

The disparate impacts of *D. simulans* and *D. melanogaster* Cuff on gene expression are potentially explained by the former’s sequestration of other *D. melanogaster* proteins into nonfunctional complexes, including the conserved transcriptional regulator C-terminal Binding Protein (CtBP; Parhad et al. 2020). While physical interaction between Cuff and CtBP is required to suppress mRNA transcription at piRNA clusters, sequestration of CtBP could affect its function in regulating protein coding genes (Figure 1A; Fang et al. 2006; Phippen et al. 2000). To evaluate this possibility, I compared gene expression changes resulting from ovarian *CtBP* knock-down (Parhad et al. 2020; <u>TableS4</u>), to those arising from *D. simulans cuff*. For *CtBP-KD1* (*P{KK108401}VIE-260B*), upregulated genes were unrelated to those upregulated by *D. simulans cuff* as compared to *D. melanogaster cuff* (Fisher’s Exact Test P-value = 0.522, Figure 5E). However, for *CtBP*-KD2 (*P{GD4268}v37609*) upregulated genes are highly significantly enriched for those upregulated by *D. simulans cuff*, with 41 genes commonly upregulated in both genotypes (***X***^2^ = 451.05, df = 1, P-value < 10^-15^, Figure 5F). Interestingly, *CtBP*-KD1 has a stronger impact on *CtBP* expression than *CtBP*-KD2 (74% as compared to 30% decrease in expression (Parhad et al. 2020)), suggesting that *D. simulans cuff* is more similar to a mild reduction in *CtBP* function. Similarly, although only 18 genes were downregulated by *D. simulans cuff*, these were significantly enriched for genes downregulated by both *CtBP* knock-downs (*CtBP*-KD1: ***X***^2^ = 188.15, df = 1, P-value < 10^-15^, *CtBP*-KD2: Fisher’s Exact Test P-value = 0.006641, Figure 5G-H). Thus, the considerable impact of *D. simulans cuff* on host gene regulation is at least partially explained by its sequestration of CtBP. Genes regulated by *D. simulans cuff* but not *CtBP* could be targets of other sequestered proteins, such as TRF2 (Parhad et al. 2020).

## DISCUSSION

Here I examined the potential for genomic autoimmunity to shape interspecific divergence in the RDC complex: a key regulator of piRNA precursor transcription. For both Rhi and Cuff I observed expanded regulation of *D. melanogaster* genes by a *D. simulans* protein or domain, consistent with the autoimmunity model. However, my observations also make two important and contrasting additions to our current understanding of genomic autoimmunity. In the case of Rhi, expanded autoimmunity of the *D. simulans* hinge domain occurs through the expected mechanism of occupancy and negative regulation of host genes. However, I demonstrated that it can be masked in an interspecific complementation assay by an incompatibility in another protein domain (chromo shadow). By contrast in the case of Cuff, I demonstrated that autoimmunity unexpectedly leads to positive regulation of host genes, most likely through the disrupted function of genic transcriptional regulators.

The increased autoimmunity of the *D. simulans* hinge domain of Rhi occurs exclusively at replication dependent histones. While Rhi has not previously been reported to regulate histone transcripts, I discovered that both the *D. melanogaster* and *D. simulans* proteins, as well as all of their fusion proteins, localize to the histone gene cluster (Figure 4C). Furthermore, the native *D. melanogaster* protein exhibits modest negative regulation of *his1* and *his2A* (Figure 4B). The histone array is therefore a clear target of Rhi, with expanded regulatory effects associated with both the hinge and chromo domains of *D. simulans*. Taken together, these observations are consistent with a model in which selection has acted on the same domains in *D. melanogaster* to reduce negative regulation of histones.

While the mechanism of Rhi recruitment to the histone gene cluster is unclear, replication-dependent histones employ a non-canonical transcriptional program that shares some features with piRNA transcription. The transcription of both piRNA precursors and *his1* is initiated by TRF2, although in the former case TRF2 is recruited by Rhi and Del rather than direct binding to the promoter (Isogai et al. 2007; Andersen et al. 2017). Rhino’s association with TRF2 may therefore recruit it indirectly to *his1* promoters. Consistent with this model, the largest Rhi peak for all transgenes occurs upstream of *his1*, and *his1* is also the most strongly negatively regulated histone by the *D. melanogaster* transgene and the chromo and hinge fusion constructs (Figure 4A). However, because the association between TRF2 and Rhi is thought to be mediated through Del, TRF2’s association cannot explain the recruitment of the *D. simulans* and chromo shadow fusion Rhi proteins, which cannot bind Del but still occupy the histone gene cluster. An alternative mechanism for Rhi recruitment is the histone modification H3K9me3, which is bound by Rhi to initiate piRNA precursor transcription (Mohn et al. 2014; Le Thomas et al. 2014). The histone methylatransferase Su(var)3-9 localizes to the histone gene cluster in salivary glands, and mutants exhibit increased histone expression (Ner et al. 2002). Given the role of Su(var)3-9 in depositing H3K9me3 during oogenesis (Yoon et al. 2008), it seems likely that this heterochromatic mark also occurs at the histone gene cluster in ovaries, potentially leading to Rhi recruitment. Regardless of the underlying mechanisms, my observations suggest that shared regulatory machinery between piRNA clusters and host genes, including both transcription factors and histone modifications, may be an important source of genomic autoimmunity.

My observations with *cuff* similarly implicate shared cofactors in the autoimmunity of nuclear piRNA proteins. *Drosophila simulans cuff* regulates the expression of hundreds of genes when compared to its *D. melanogaster* counterpart, most likely through the sequestration of transcriptional regulators like CtBP. Enhanced affinity for co-factors such as CtBP might have facilitated expanded TE silencing in the *D. simulans* lineage, with downstream impacts on host genes having been ameliorated by regulatory changes elsewhere in the system. This is broadly consistent with the proposed selection regime in the autoimmunity model, in which invading or escaping TEs select for expanded piRNA-mediated regulation, and subsequently compensatory mutations arise that decrease off target effects (Blumenstiel et al. 2016).

## MATERIALS AND METHODS

### Data sets and quality control

*rhi* and *del* ovarian RNA-seq, small RNA-seq and ChIP seq (*rhi* only) data sets are from Parhad *et al.* (2017). *cuff* and *CtBP* ovarian RNA-seq data were from Parhad *et al.* (2020). All Illumina libraries downloaded and analyzed are described in Table S5. Data were downloaded from the NCBI Sequenced Read Archive. Adaptors were removed and low-quality bases were trimmed from all raw-reads using trim-galore (Krueger 2015).

### RNA-seq analysis

RNA-seq reads were aligned to release 6.33 of the *D. melanogaster* transcriptome using Kallisto (Bray et al. 2016), in order to estimate the abundance of each transcript. The estimated number of reads was then summed across all transcripts from the same gene to obtain the estimated read count for each gene. For histone gene copies, the estimated number of reads were further summed across all copies of the gene, since reads cannot be reliably assigned to individual copies. Genic read counts were then used to estimate differential expression.

For *del* and *cuff* mutants and transgenic rescues, as well as for *CtBP* knock-down and control flies (*white* knock-down) only one biological replicate was available for each genotype. We therefore used DESeq to estimate differential expression (method=“blind”,sharingMode=“fit-only” (Anders & Huber 2010), and significant differences were detected using a negative binomial test. Genes with fewer than 50 reads in all samples were excluded. For *rhi* mutants and transgenic rescues, two biological replicates were available for each genotype. We therefore estimated differential expression and detected statistical significance with DEseq2 (Love et al. 2014). Genes with fewer than 50 average reads across samples were excluded. Regardless of the analysis package, a gene was considered differentially expressed if the adjusted P-value was less than 0.05.

### ChIP-seq analysis

Chip Seq reads were aligned to a reference genome using BWA (Li & Durbin 2009). To avoid complications of multiply-mapping reads in the histone gene cluster, reads were aligned to a custom version of the dm6 reference genome in which the histone array (2L:21,403,672-21,543,688) was replaced with a single representative copy of the histone repeat (2L:21,482,367-21,487,518), as in McKay *et al.* (2015). Peaks of Rhi occupancy were detected using MACS2 with broad-peaks settings at a significance cutoff of 0.1 (Zhang et al. 2008; Feng et al. 2012). A gene was considered occupied by Rhi if a peak occurred within 1000 nt of a transcription start site, or anywhere in the transcript body inclusive of introns, based on flybase annotated transcripts.

To generate sliding window analyses of Rhi occupancy of the histone gene cluster, I first extracted read alignments overlapping the cluster via samtools (Li et al. 2009). I then calculated the nucleotide coverage, normalized to the number of aligned sequencing reads, using bedtools genomecov (Quinlan and Hall 2010). Sliding window estimates of mean coverage, relative to input were then calculated using the rollapply function from the zoo package (Zeileis & Grothendieck 2005) in R version 3.6.1 (R Development Core Team 2008).

### small-RNA analysis

Adapters were trimmed from small RNAs and putative miRNAs (18-22nt) and piRNAs (23-32 nt) were identified using trim galore (Krueger 2015). Putative miRNAs were then aligned to all annotated miRNAs in the (dm6) reference assembly, whereas piRNAs were aligned to the custom reference with a single copy of the histone array. Sliding window analyses of piRNA abundance across the histone gene cluster was performed as with the ChIP-seq data except the coverage was normalized to number of reads aligning to miRNAs from the same library. Similarly, differential piRNA abundance of individual histone genes was generated by counting the number of reads overlapping each annotated transcript using samtools (Li & Durbin 2009), and normalizing to the number of reads aligning to miRNAs from the same library.

### Statistics and Data Visualization

Statistical testing was performed in R version 3.6.1 (R Development Core Team 2008). Data were wrangled using tidyverse (Wickham et al. 2019), and represented using UpSetR (Conway et al. 2017), and ggplot2 (Wickham 2011).

## REFERENCES

Andersen PR, Tirian L, Vunjak M, Brennecke J. 2017. A heterochromatin-dependent transcription machinery drives piRNA expression. Nature. 549:54–59.

Anders S, Huber W. 2010. Differential expression analysis for sequence count data. Genome Biol. 11:R106.

Aruna S, Flores HA, Barbash DA. 2009. Reduced fertility of Drosophila melanogaster hybrid male rescue (Hmr) mutant females is partially complemented by Hmr orthologs from sibling species. Genetics. 181:1437–1450.

Blumenstiel JP, Erwin AA, Hemmer LW. 2016. What Drives Positive Selection in the Drosophila piRNA Machinery? The Genomic Autoimmunity Hypothesis. Yale J. Biol. Med. 89:499–512.

Brand CL, Cattani MV, Kingan SB, Landeen EL, Presgraves DC. 2018. Molecular Evolution at a Meiosis Gene Mediates Species Differences in the Rate and Patterning of Recombination. Curr. Biol. 28:1289–1295.e4.

Bray N, Pimentel H, Melsted P, Pachter L. 2016. Near-optimal RNA-Seq quantification with kallisto. Nat. Biotechnol. 34:525–527.

Chen Y-CA et al. 2016. Cutoff Suppresses RNA Polymerase II Termination to Ensure Expression of piRNA Precursors. Mol. Cell. 63:97–109.

Conway JR, Lex A, Gehlenborg N. 2017. UpSetR: an R package for the visualization of intersecting sets and their properties. Bioinformatics. 33:2938–2940.

Crysnanto D, Obbard DJ. 2019. Widespread gene duplication and adaptive evolution in the RNA interference pathways of the Drosophila obscura group. BMC Evol. Biol. 19:99.

Czech B et al. 2018. piRNA-Guided Genome Defense: From Biogenesis to Silencing. Annu. Rev. Genet. 52:131–157.

El Baidouri M, Panaud O. 2013. Comparative genomic paleontology across plant kingdom reveals the dynamics of TE-driven genome evolution. Genome Biol. Evol. 5:954–965.

Fang M et al. 2006. C-terminal-binding protein directly activates and represses Wnt transcriptional targets in Drosophila. EMBO J. 25:2735–2745.

Feng J, Liu T, Qin B, Zhang Y, Liu XS. 2012. Identifying ChIP-seq enrichment using MACS. Nat. Protoc. 7:1728–1740.

Flores HA, Bubnell JE, Aquadro CF, Barbash DA. 2015. The Drosophila bag of marbles Gene Interacts Genetically with Wolbachia and Shows Female-Specific Effects of Divergence Malik, HS, editor. PLoS Genet. 11:e1005453.

Gilbert C, Schaack S, Pace JK, Brindley PJ, Feschotte C. 2010. A role for host-parasite interactions in the horizontal transfer of transposons across phyla. Nature. 464:1347–1350.

Isogai Y, Keles S, Prestel M, Hochheimer A, Tjian R. 2007. Transcription of histone gene cluster by differential core-promoter factors. Genes Dev. 21:2936–2949.

Kidwell MG. 1983. Evolution of hybrid dysgenesis determinants in Drosophila melanogaster. Proc. Natl. Acad. Sci. U. S. A. 80:1655–1659.

Kolaczkowski B, Hupalo DN, Kern AD. 2011. Recurrent adaptation in RNA interference genes across the Drosophila phylogeny. Mol. Biol. Evol. 28:1033–1042.

Krueger F. 2015. Trim galore. A wrapper tool around Cutadapt and FastQC to consistently apply quality and adapter trimming to FastQ files. 516:517.

Le Thomas A et al. 2014. Transgenerationally inherited piRNAs trigger piRNA biogenesis by changing the chromatin of piRNA clusters and inducing precursor processing. Genes Dev. 28:1667–1680.

Lewis SH, Salmela H, Obbard DJ. 2016. Duplication and diversification of Dipteran Argonaute genes, and the evolutionary divergence of Piwi and Aubergine. Genome Biol. Evol. doi: 10.1093/gbe/evw018.

Li H et al. 2009. The Sequence Alignment/Map format and SAMtools. Bioinformatics. 25:2078–2079.

Li H, Durbin R. 2009. Fast and accurate short read alignment with Burrows-Wheeler transform. Bioinformatics. 25:1754–1760.

Love MI, Huber W, Anders S. 2014. Moderated estimation of fold change and dispersion for RNA-seq data with DESeq2. Genome Biol. 15:550.

McKay DJ et al. 2015. Interrogating the function of metazoan histones using engineered gene clusters. Dev. Cell. 32:373–386.

Mohn F, Sienski G, Handler D, Brennecke J. 2014. The rhino-deadlock-cutoff complex licenses noncanonical transcription of dual-strand piRNA clusters in Drosophila. Cell. 157:1364–1379.

Naito K et al. 2006. Dramatic amplification of a rice transposable element during recent domestication. Proc. Natl. Acad. Sci. U. S. A. 103:17620–17625.

Ner SS, Harrington MJ, Grigliatti TA. 2002. A role for the Drosophila SU(VAR)3-9 protein in chromatin organization at the histone gene cluster and in suppression of position-effect variegation. Genetics. 162:1763–1774.

Obbard DJ, Gordon KHJ, Buck AH, Jiggins FM. 2009. The evolution of RNAi as a defence against viruses and transposable elements. Philos. Trans. R. Soc. Lond. B Biol. Sci. 364:99–115.

Ozata DM, Gainetdinov I, Zoch A, O’Carroll D, Zamore PD. 2019. PIWI-interacting RNAs: small RNAs with big functions. Nat. Rev. Genet. 20:89–108.

Palmer WH, Hadfield JD, Obbard DJ. 2018. RNA-Interference Pathways Display High Rates of Adaptive Protein Evolution in Multiple Invertebrates. Genetics. 208:1585–1599.

Parhad SS et al. 2020. Adaptive Evolution Targets a piRNA Precursor Transcription Network. Cell Reports. 30:2672–2685.e5. doi: 10.1016/j.celrep.2020.01.109.

Parhad SS, Tu S, Weng Z, Theurkauf WE. 2017. Adaptive Evolution Leads to Cross-Species Incompatibility in the piRNA Transposon Silencing Machinery. Dev. Cell. 43:60–70.e5.

Phippen TM et al. 2000. Drosophila C-terminal binding protein functions as a context-dependent transcriptional co-factor and interferes with both mad and groucho transcriptional repression. J. Biol. Chem. 275:37628–37637.

Quinlan AR, Hall IM. 2010. BEDTools: a flexible suite of utilities for comparing genomic features. Bioinformatics. 26:841–842.

Quinlan AR, Hall IM. BEDTools: a flexible suite of utilities for comparing genomic features. Bioinformatics [Internet]. 2010; 26: 841–2.

R Development Core Team. 2008. R: A Language and Environment for Statistical Computing. http://www.r-project.org.

Reiss D et al. 2019. Global survey of mobile DNA horizontal transfer in arthropods reveals Lepidoptera as a prime hotspot. PLoS Genet. 15:e1007965.

Senti K-A, Brennecke J. 2010. The piRNA pathway: a fly’s perspective on the guardian of the genome. Trends Genet. 26:499–509.

Simkin A, Wong A, Poh Y-P, Theurkauf WE, Jensen JD. 2013. Recurrent and recent selective sweeps in the piRNA pathway. Evolution. 67:1081–1090.

Smothers JF, Henikoff S. 2001. The hinge and chromo shadow domain impart distinct targeting of HP1-like proteins. Mol. Cell. Biol. 21:2555–2569.

Vermaak D, Henikoff S, Malik HS. 2005. Positive selection drives the evolution of rhino, a member of the heterochromatin protein 1 family in Drosophila. PLoS Genet. 1:96–108.

Vermaak D, Malik HS. 2009. Multiple roles for heterochromatin protein 1 genes in Drosophila. Annu. Rev. Genet. 43:467–492.

Wang L, Barbash DA, Kelleher ES. 2020. Adaptive evolution among cytoplasmic piRNA proteins leads to decreased genomic auto-immunity. PLOS Genetics. 16:e1008861. doi: 10.1371/journal.pgen.1008861.

Wickham H. 2011. ggplot2. WIREs Comp Stat. 3:180–185.

Wickham H et al. 2019. Welcome to the Tidyverse. Journal of Open Source Software. 4:1686.

Yang H-P, Barbash DA. 2008. Abundant and species-specific DINE-1 transposable elements in 12 Drosophila genomes. Genome Biol. 9:R39.

Yi M et al. 2014. Rapid evolution of piRNA pathway in the teleost fish: implication for an adaptation to transposon diversity. Genome Biol. Evol. 6:1393–1407.

Yoon J et al. 2008. dSETDB1 and SU(VAR)3-9 sequentially function during germline-stem cell differentiation in Drosophila melanogaster. PLoS One. 3:e2234.

Yu B et al. 2018. Structural insights into Rhino-Deadlock complex for germline piRNA cluster specification. EMBO Rep. 19. doi: 10.15252/embr.201745418.

Yu B et al. 2015. Structural insights into Rhino-mediated germline piRNA cluster formation. Cell Res. 25:525–528.

Zeileis A, Grothendieck G. 2005. zoo: S3 Infrastructure for Regular and Irregular Time Series. Journal of Statistical Software, Articles. 14:1–27.

Zhang Y et al. 2008. Model-based analysis of ChIP-Seq (MACS). Genome Biol. 9:R137.

Zhang Z et al. 2014. The HP1 homolog rhino anchors a nuclear complex that suppresses piRNA precursor splicing. Cell. 157:1353–1363.

